# Deciphering the HDAC6-Mediated Regulation of MLLT3 in Myeloid Progenitor Cell Fate: Insights into Stem Cell Differentiation Dynamics

**DOI:** 10.64898/2026.04.30.721939

**Authors:** Mabu P Subahan, Anil Aribandi, Arunasree M Kalle

## Abstract

Mixed-lineage leukemia translocated to 3 (MLLT3) is vital for maintaining the stemness of hematopoietic stem cells. Loss of MLLT3 in megakaryocyte (MK)-erythrocyte progenitor (MEP) cells leads to its differentiation into MKs. Despite its significance in stemness, the regulatory mechanism of MLLT3 during differentiation remains elusive. In this study, we investigate the regulatory role of histone deacetylase 6 (HDAC6) in modulating MLLT3 levels *via* heat shock protein 90 (Hsp90) activation during myeloid lineage differentiation into MKs, monocytes, and macrophages. We found that HDAC6 activates Hsp90 through deacetylation, enabling Hsp90 to retain MLLT3 in the cytoplasm where protein kinase C (PKC) phosphorylates MLLT3 at serine residues; leading to loss of MLLT3 during MK and macrophage differentiation but not during monocyte differentiation. This research provides valuable insights into the regulatory mechanisms underlying myeloid lineage commitment and opens new avenues for future investigations into stem cell biology and therapeutic applications.

## Introduction

Self-renewal is the ability of stem cells and progenitor cells to proliferate while maintaining their multipotency. This fundamental characteristic is crucial for tissue homeostasis and regeneration^1^. In the hematopoietic system, progenitor cells derived from hematopoietic stem cells (HSCs) such as the common lymphoid progenitor, common myeloid progenitor, and megakaryocyte (MK)- erythrocyte progenitor (MEP) possess short-term self-renewal capacity until they receive signals that trigger their differentiation^2^.

Rather than conferring, mixed-lineage leukemia translocated to 3 (MLLT3) has gained significant attention as the key factor involved in maintaining stemness to HSCs^3^. MLLT3 functions as an essential component of the super-elongation complex (SEC). SEC plays a crucial role in enhancing RNA polymerase II transcriptional elongation by suppressing transient polymerase pausing^4–6^. By recognizing histone H3 crotonylated and acetylated lysines through its YEATS domain, MLLT3 facilitates SEC chromatin binding^7,8^. MLLT3 has not only been previously identified as a fusion partner of MLL genes^9,10^,its critical role in HSCs maintenance and differentiation has also been clearly established^3^. A study by Pina et al.^11^ demonstrated that MLLT3 is a critical regulator of differentiation of erythroid and MK from MEP cells. It is known that the loss of MLLT3 transcriptional activity promotes differentiation of the progenitor cells; however, its role in differentiation into other blood cell lineages is not yet clearly understood.

Heat shock protein 90 (Hsp90) is a highly conserved and abundantly expressed chaperone protein. Hsp90 is known for its role in protein folding and maintenance of protein stability^12^. Recent studies have unveiled its involvement in the nucleo-cytoplasmic compartmentalization of target proteins thereby regulating their transcriptional activity^13^. MLLT3 has also been identified as one of the client proteins of Hsp90^14^. Hsp90 itself undergoes post-translational modifications including acetylation which can modulate its binding affinity to its client proteins. While hyperacetylation of Hsp90 reduces its affinity, deacetylation by histone deacetylase 6 (HDAC6) enhances its chaperone activity^15^. Hsp90 inhibitors are identified as potential candidates for the treatment of acute myeloid leukemia (AML), suggesting its role in HSC differentiation.

In a study by Messaoudi et al^16^., HDAC6’s essential role in human MK differentiation from MEP cells was discovered. Given the involvement of HDAC6 in MK differentiation and its impact on Hsp90 function (which in turn regulates MLLT3 transcriptional activity to maintain stemness in HSC), we hypothesized a potential indirect negative relationship between MLLT3 and HDAC6 during MEP differentiation into MKs.

In this study, we present compelling evidence confirming the negative regulation of MLLT3 by HDAC6 *via* Hsp90. Additionally, we demonstrate the significance of PKC-mediated serine phosphorylation which leads to the loss of MLLT3 during terminal myeloid lineage differentiation into MKs and macrophages, but not during intermediate cell differentiation into monocytes.

Our findings provide insights into the intricate regulatory network governing myeloid lineage differentiation and highlight the critical roles of MLLT3, Hsp90, and HDAC6 in this process.

## Methods

### Experimental Models used in the study

Human chronic myelogenous leukemia cell line K562, human monocytic leukemia cell lines, THP1, human promyelocytic leukemia cell lines, HL60 and human hematopoietic stem cells CD34+ cells were used as model systems in the study. K562 cells when induced with PMA differentiate into megakaryocyte. THP1 cells when treated with PMA for 2 days, differentiate into macrophage. HL60 cells when induced with PMA for 2 days differentiate into monocyte and when induced with PMA for 5 days differentiate into macrophages. CD34+ cells under various factors can be differentiated into variety of blood cells. TPO induces MK differentiation, StemSpan™ Myeloid Expansion Supplement II (100X) leads to monocyte differentiation which when grown further in ImmunoCult™ – SF Macrophage Medium differentiate into macrophages.

### Cell culture

All the cells (K562, THP1 and HL-60) were procured from the National Center for Cell Science cell repository, India. K562, THP1 and HL-60 cells were cultured in RPMI 1640 medium supplemented with 10% FBS and 1× *Penicillin* and *Streptomycin* and grown at 37°C and 5% CO_2_.

The K562 cells were subjected to MK differentiation, and THP1 and HL-60 were subjected to macrophage differentiation using phorbol-myristate-12 acetate (PMA) treatment for 24 and 48 hours. The exponentially growing cells were seeded in 6-well plates and subjected to different inhibitor treatments–5 mM Tubastatin A (TUB, HDAC6 inhibitor), 10 mM Sotrastaurin (SOT, protein kinase C inhibitor) and 5 mM Novobiocin (NOV, Hsp90 inhibitor) for 24 and 48 hours. For combination, the cells were treated with inhibitor first, and were given PMA one hour later.

### CD34^+^ cell culture

CD34^+^ cells were purchased from StemSpan Technologies. The cells were cultured in phenol red-free IMDM supplemented with 10 mg/mL of FBS, 10 μg/mL of bovine insulin, 200 μg/mL of human transferrin, 40 μg/mL of human low-density lipoprotein (Sigma), 2 mmol/L L-glutamine (Sigma), 10^−4^ mol/L 2-mercaptoethanol (Sigma), and growth factors as per manufacturer protocol, and grown at 37°C and 5% CO_2_.

Thrombopoietin (TPO) treatment for 14 days induced MK differentiation of the CD34^+^ cells. The cells were assayed for MK differentiation after treatment with TPO for 14 days, followed by labeling with FITC-labeled anti-CD41 antibody and analysis using fluorescence-activated cell sorting with appropriate gating and cell sorting. StemSpan™ Myeloid Expansion Supplement II (100×) induced monocyte differentiation of CD34^+^ cells following 14 days of treatment and were analyzed using FITC-labeled anti-CD64 antibody. The monocytic cells were treated with ImmunoCult-SF Macrophage Medium to promote macrophage production following 8 days of treatment from CD64^+^ monocytes. The differentiation was evaluated using fluorescein isothiocyanate-labeled anti-CD68 antibody.

The exponentially growing cells were seeded in 6-well plates and subjected to different inhibitor treatments–5 mM TUB (HDAC6 inhibitor), 10 mM SOT (Protein Kinase C inhibitor) and 5 mM NOV (Hsp90 inhibitor) where the cells were treated with inhibitors first, followed by PMA addition one hour later.

### RNA isolation and qPCR

Total RNA was extracted by TRI reagent (Sigma) according to the study conducted by Doddi et al^17^. The RNA concentration was estimated using Nanodrop. For cDNA synthesis, iScript™ cDNA Synthesis Kit was used according to the manufacturer’s protocol. The amplification of target genes of the cDNA synthesized was carried out with gene specific primers using KAPA SYBR FAST qPCR Master Mix (2×) in Realtime PCR machine (Applied Biosystems). The expression levels were calculated after normalizing with the control cells using the 2^−ΔΔCT^ method.

### Immunoblot analysis

The total cell lysates, cytoplasmic and nuclear fractions of the (K562, THP1, HL-60 and CD34^+^ cells) were prepared using radioimmunoprecipitation assay lysis buffer consisting of protease inhibitors and phosphatase inhibitors as per the protocol described earlier^18^. The protein concentration in the lysate and subcellular fractions was estimated by Bradford methods and stored at −20°C until further use. A 40 to 60 µg of the protein was separated on 10% to 12% SDS-PAGE followed by Western blotting as per standard protocol. The protein bands were developed and visualized using ChemiDoc MP imaging system (Bio-Rad).

### Immunofluorescence

The cells (K562, THP1 and HL-60) were fixed with 4% paraformaldehyde for 15 minutes in dark. The cells were washed twice with 1× PBS and permeabilized with 0.5% Triton X-100 prepared in PBS for 30 minutes at room temperature. The cells were washed thrice with PBST (phosphate-buffered saline with Tween 20) and blocked with blocking buffer (3% BSA in PBST and 0.1% Triton X-100) for one hour at room temperature. This was followed by five washed using PBS. The cells were then incubated with 100 µl primary antibody prepared in 3% BSA in PBST (1:100) overnight at 4°C. The cells were washed five times with PBS, followed by incubation with secondary antibody prepared in PBST (1:500) for 2 hours at room temperature in dark. The cells were washed five times with PBS, mounted on cover slip using DPX Mountant and observed under confocal microscope (Leica–microscope brand and model).

### Flow cytometer

The differentiation of cells into MKs (K562, CD34^+^ cells), macrophages (THP1, HL-60, CD34^+^ cells) and monocytes (HL-60, CD34^+^ cells) was evaluated by BD FACS Aria flowcytomer using FITC-labeled surface markers (CD41-MKs, CD64-monocytes and CD68-macrophages) as per the protocol given by the manufacturer (BD Biosciences).

### Statistical analysis

The standard deviation of the data obtained from three independent experiments with duplicates denotes the results. Statistical analysis of the experiments was performed on the GraphPad Prism 10.0.1 software using the one-way ANOVA test with p-values indicating the significance of the data *p-value < 0.05, **p-value < 0.01 and ***p-value < 0.001.

## Results

### HDAC6 regulates MLLT3 nuclear localization via Hsp90 during MK differentiation

Given the interaction between HDAC6 and Hsp90 and their impact on MLLT3 regulation, we sought to investigate their involvement during MK differentiation. We used Tubastatin (TUB) which inhibits HDAC6 function and novobiocin (NOV) which inhibits Hsp90 function and tried to evaluate their regulatory role on MLLT3 during MK differentiation. We used PMA-induced K562 cells and TPO-induced CD34+ cells as model systems (Figure 1A). Upon induction with TPO or PMA alone, we observed a significant loss of MLLT3 protein levels and a concurrent increase in HDAC6 protein levels relative to the untreated controls. To investigate the regulatory mechanisms governing MLLT3 stability, cells were pre-treated with TUB or NOV prior to the induction of differentiation. Interestingly, pre-treatment with either inhibitor reversed the trends observed during standard differentiation. Under both the TUB and NOV pre-treatment conditions, MLLT3 protein levels were notably higher, while HDAC6 protein levels were decreased, when compared directly to the TPO/PMA differentiation conditions. (Figure 1B). During MK differentiation, we observed a reduction in acetylated Hsp90 levels (Figure 1B), indicative of an active Hsp90 state, which drives the subsequent decrease in MLLT3 protein levels. (Figure 1B). Conversely, targeted inhibition with either TUB or NOV promoted Hsp90 hyperacetylation. (Figure 1B). This shift to an inactive, hyperacetylated Hsp90 state disrupts the MLLT3 clearance mechanism that is typically engaged during MK differentiation, ultimately leading to the restoration of MLLT3 expression. (Figure 1B). Together, these findings demonstrate that MLLT3 degradation during megakaryopoiesis is tightly regulated by the HDAC6-Hsp90 axis, where HDAC6-mediated deacetylation maintains Hsp90 in an active state to facilitate MLLT3 clearance. (Figure 1B). Furthermore, the MLLT3 RNA levels also decreased significantly in PMA-treated K562 cells and TPO-treated CD34^+^ cells. The RNA levels were partially restored in NOV and TUB treated cells (Figure 1C). To determine the subcellular compartment in which the MLLT3 clearance mechanism occurs during MK differentiation, we isolated and analyzed both cytoplasmic and nuclear fractions. Immunoblot analysis clearly indicated that the differentiation-induced loss of MLLT3 occurs specifically within the nuclear compartment, while the cytoplasmic levels of MLLT3 showed no change. (Figure 1D). Furthermore, this targeted nuclear degradation was prevented in the presence of TUB and NOV which successfully restored MLLT3 levels in the nucleus. (Figure 1D).

**Figure 1.**
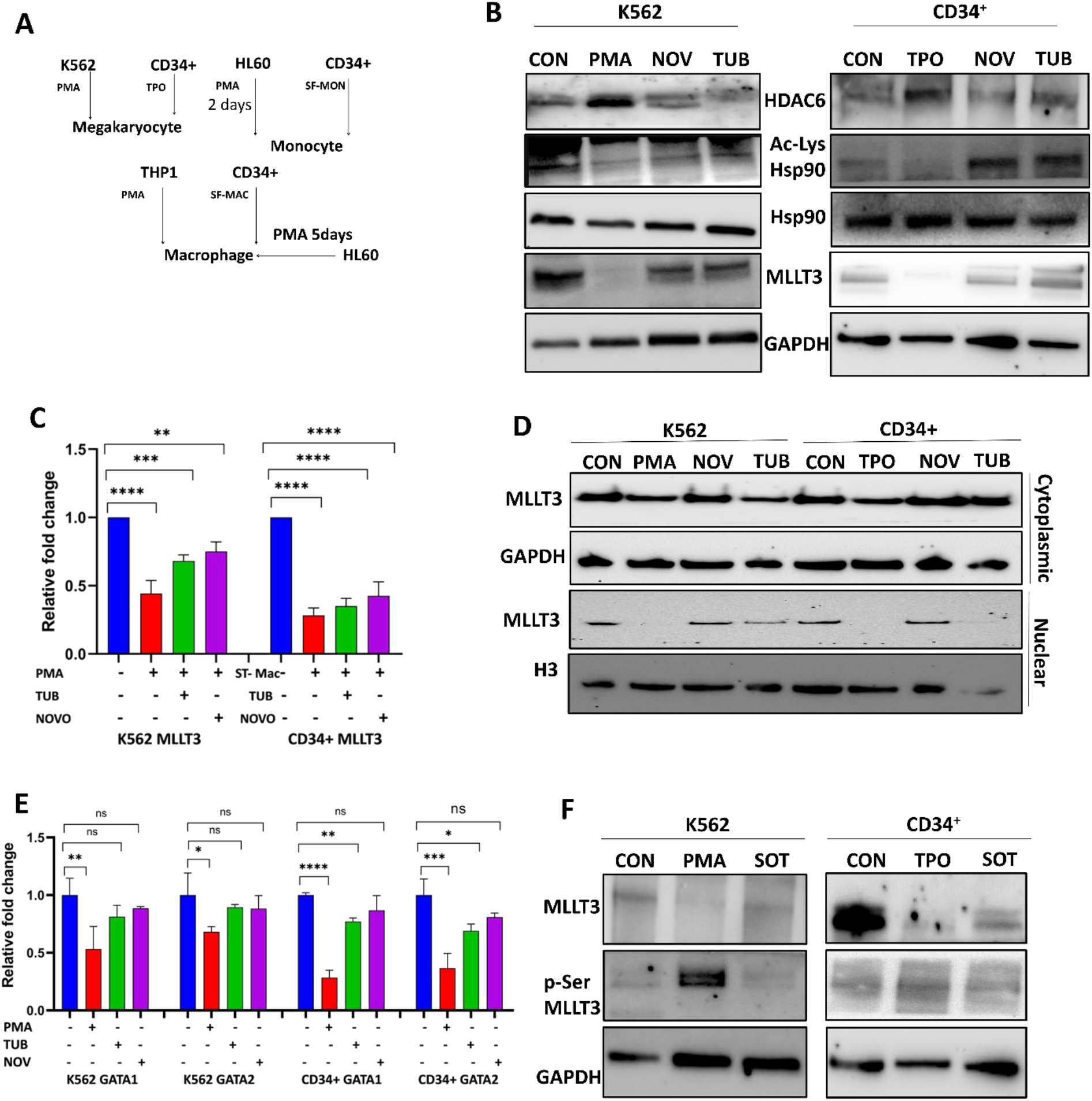
HDAC6 promotes cytoplasmic shuttling of MLLT3 via Hsp90 leading to serine phosphorylation of MLLT3 by PKC. (A) Model systems used in the study. (B) Immunoblot analysis of HDAC6, HSp90, Ac-Hsp90 and MLLT3 protein levels in presence or absence of Tubastatin (TUB) and Novobiocin (NOV) during MK differentiation of K562 cells (PMA) or CD34+ cells (TPO). (C) Real-time PCR analysis of MLLT3 mRNA levels during MK differentiation of K562 cells (PMA) or CD34+ cells (TPO). (D) Western blots showing the nuclear and cytoplasmic levels of MLLT3 in presence or absence of Tubastatin (TUB) and Novobiocin (NOV) during MK differentiation of K562 cells (PMA) or CD34+ cells (TPO). (E) Real-time PCR analysis of mRNA levels of GATA1 and GATA2 transcription factors, two of the MLLT3 transcriptional effector genes. (F) P-Serine immunoblot analysis of MLLT3 in presence of Sotrastaurin (SOT), PKC inhibitor during MK differentiation.

### Serine phosphorylation of MLLT3 by PKC results in loss of nuclear MLLT3 during MK differentiation

To understand the functional significance of MLLT3 restoration following HDAC6 and Hsp90 inhibition, we evaluated the downstream transcriptional regulation of known MLLT3 target genes. Specifically, we assessed the mRNA levels of *GATA1* and *GATA2*, which are critical factors that promote the erythroid differentiation of megakaryocyte-erythroid progenitor (MEP) cells. Quantitative analysis revealed that the mRNA expression levels of both *GATA1* and *GATA2* strongly correlated with MLLT3 protein levels across the various treatment conditions. Taken together, these data functionally validate the regulatory axis, indicating that HDAC6 exerts a negative regulatory effect on MLLT3 and its downstream transcriptional targets via Hsp90. (Figure 1E). To explore the mechanism driving MLLT3 protein loss during MK differentiation, we hypothesized that phosphorylation might be a precipitating event. Initial *in silico* analysis using kinase-specific phosphorylation prediction tools (NetPhos 3.1, PhosphoPredict, UniProt) strongly implicated the involvement of Protein Kinase C (PKC). To validate this experimentally, we evaluated the phosphorylation status of MLLT3. During standard MK differentiation, we observed a marked increase in serine-phosphorylated MLLT3 (phospho-MLLT3) levels. Crucially, treatment with the PKC inhibitor Sotrastaurin (Sot) reversed this effect, resulting in a distinct decrease in phospho-MLLT3 while concurrently restoring total MLLT3 protein levels. Together, these data strongly indicate that PKC-mediated serine phosphorylation targets MLLT3 for destabilization and subsequent clearance during megakaryopoiesis. (Figure 1F).

### Inhibition of HDAC6 or Hsp90 inhibited MK differentiation

To functionally corroborate the regulatory roles of HDAC6, Hsp90, and PKC during megakaryopoiesis, we evaluated cellular maturation using flow cytometric (FACS) analysis. In both PMA-induced K562 cells (Figure 2A), and TPO-induced primary CD34+ cells, (Figure 2B) the targeted inhibition of HDAC6 (using TUB), Hsp90 (using NOV), or PKC (using SOT) significantly impeded MK differentiation. This phenotypic block indicates that the coordinated clearance of MLLT3 by these signaling nodes is a requisite event for successful MK lineage commitment. Furthermore, these biochemical and cytometric findings were visually validated via immunofluorescence microscopy. Analysis of K562 cells undergoing MK differentiation revealed that the pharmacological inactivation of HDAC6, Hsp90, or PKC resulted in a distinct accumulation of MLLT3, as evidenced by a marked increase in MLLT3-specific red fluorescence (Figure 2C). This *in situ* observation aligns closely with our immunoblot data, confirming the robust restoration of MLLT3 protein levels when this degradation axis is disrupted. (Figure 2C).

**Figure 2.**
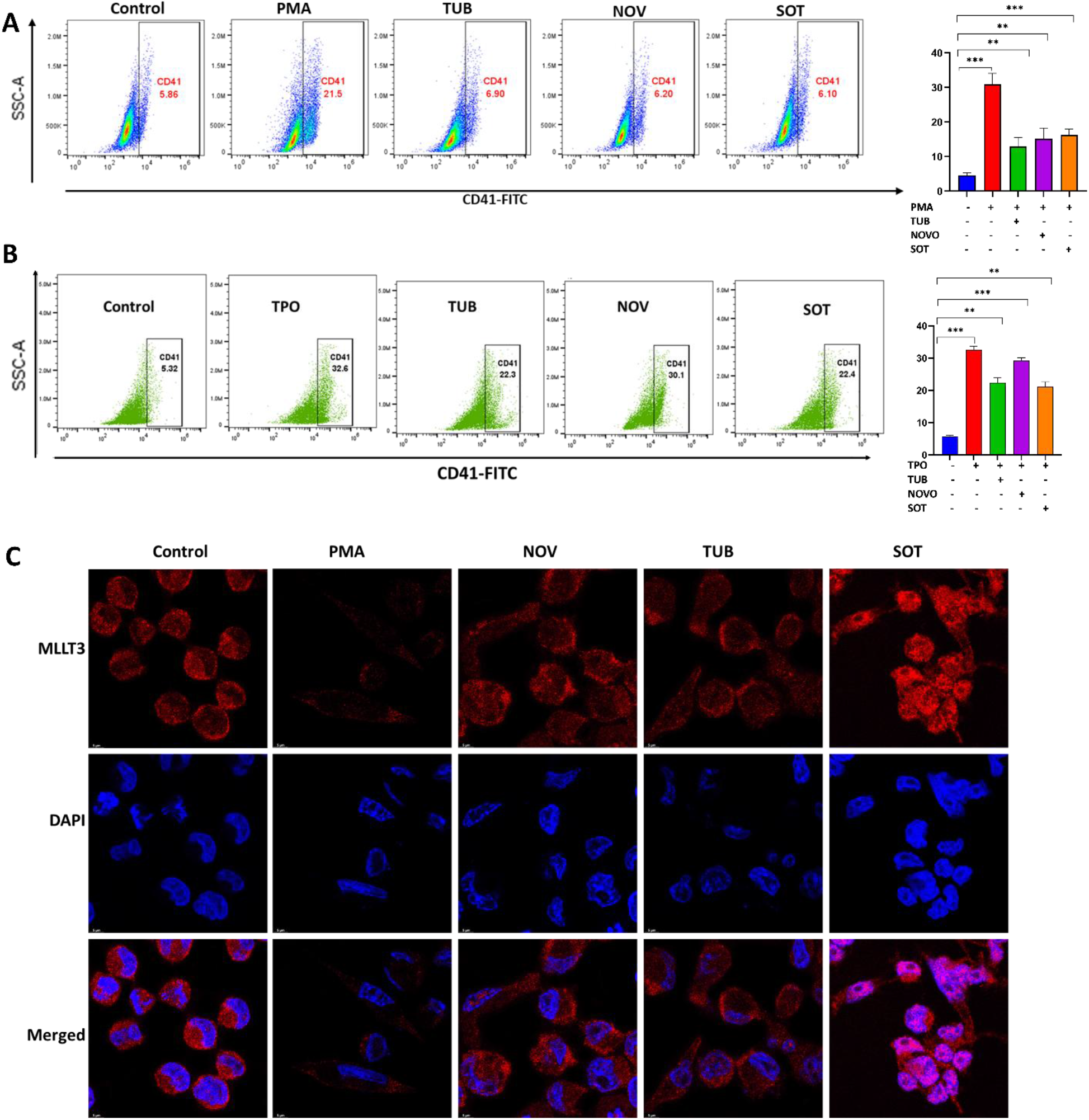
Inhibition of HDAC6, Hsp90 or PKC restores nuclear MLLT3 levels and inhibit MK differentiation. (A) FACS analysis of K562 cells in presence or absence of Tubastatin (TUB), Novobiocin (NOV) or Sotrastaurin (SOT) during MK differentiation (PMA) using FITC-CD41as MK differentiation marker. The graph below shows the % of differentiated cells. (B) FACS analysis of CD34+ cells in presence or absence of Tubastatin (TUB), Novobiocin (NOV) or Sotrastaurin (SOT) during MK differentiation (TPO) using FITC-CD41as MK differentiation marker. The graph below shows the % of differentiated cells. (C) Confocal microscopy of K562 cells undergoing MK differentiation (PMA) in presence or absence of Tubastatin (TUB), Novobiocin (NOV) or Sotrastaurin (SOT) showing MLLT3 (Red) distribution. Blue fluorescence (DAPI) indicates nucleus.

### The PKC-HDAC6-Hsp90 regulatory axis drives MLLT3 clearance during macrophage differentiation

To investigate whether the targeted degradation of MLLT3 is a conserved mechanism across other myeloid lineages, we evaluated its fate during macrophage differentiation. For these experiments, we utilized PMA-induced THP1 cells and macrophage colony-stimulating factor (M-CSF)- induced primary CD34+ cells.

Consistent with our observations in the megakaryocytic lineage, the induction of macrophage differentiation resulted in a pronounced decrease in total MLLT3 protein levels. (Figure 3A) This clearance was accompanied by an increase in HDAC6 expression, a reduction in Hsp90 acetylation, and enhanced MLLT3 serine phosphorylation (phospho-MLLT3). (Figure 3A) Crucially, targeted pharmacological intervention using the specific inhibitors TUB, NOV, or SOT successfully rescued total MLLT3 levels. (Figure 3A). Mechanistically, inhibition of this pathway reduced phospho-MLLT3 accumulation and restored Hsp90 to a hyperacetylated, inactive state, confirming that the degradation machinery is identical to that seen in MK cells. Furthermore, FACS analysis demonstrated that treatment with TUB, NOV, or SOT effectively blocked macrophage differentiation, (Figure 3B) mirroring the phenotypic arrest seen in our MK models. These biochemical and functional findings were visually corroborated by immunofluorescence imaging. (Figure 3C). Together, these data indicate that the PKC-HDAC6-Hsp90 regulatory axis governing MLLT3 phosphorylation and subsequent clearance is a shared and necessary event for proper myeloid differentiation into both megakaryocytes and macrophages.

**Figure 3.**
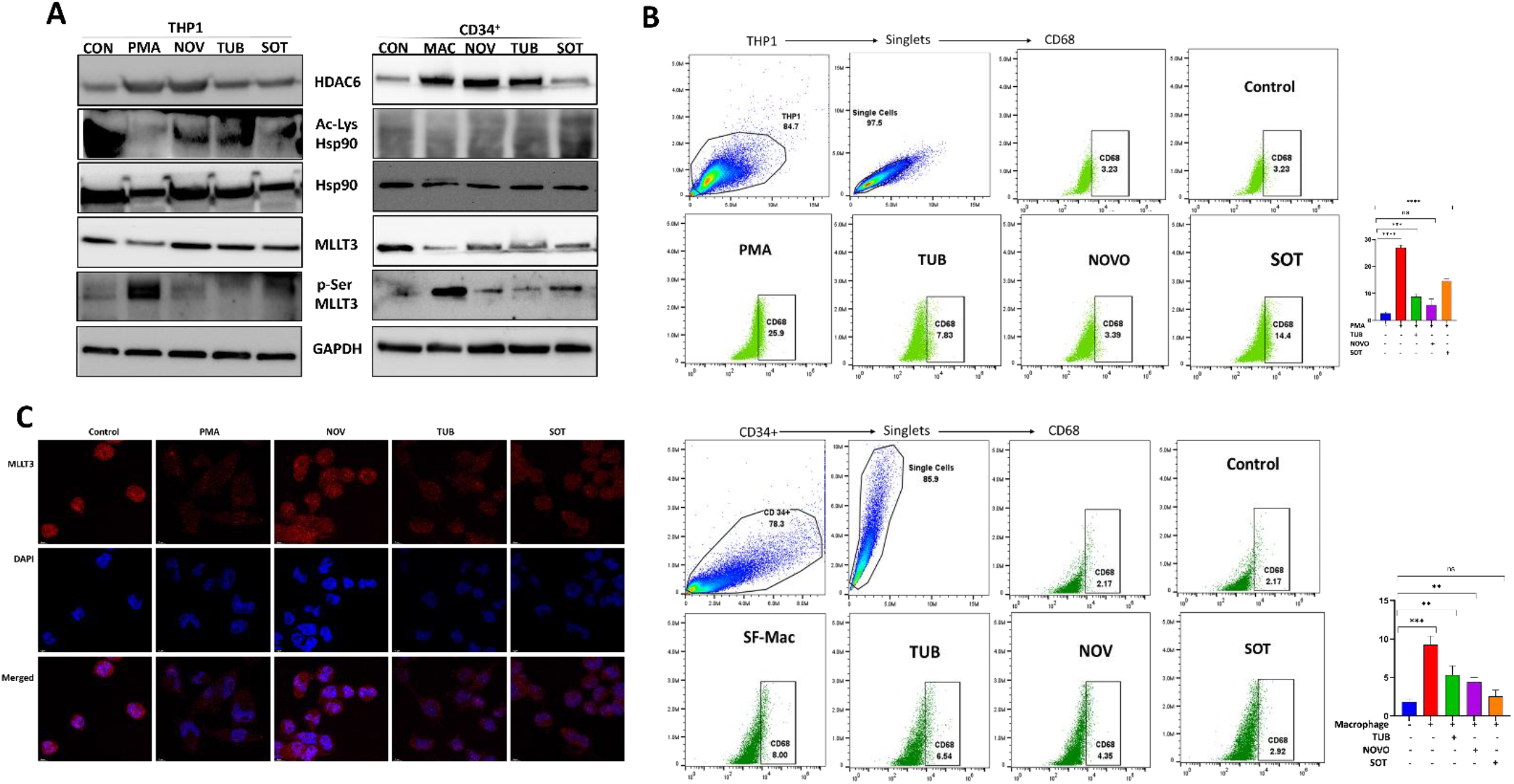
MLLT3 is translocated to cytoplasm by HDAC6-activated Hsp90 and is phosphorylated by PKC during macrophage differentiation. (A) Immunoblot analysis of HDAC6, HSp90, Ac-Hsp90, MLLT3 and p-Ser-MLLT3 protein levels in presence or absence of Tubastatin (TUB), Novobiocin (NOV) and Sotrastaurin (SOT) during macrophage differentiation of THP1 cells (PMA) and CD34+ cells (SF-Mac). A. FACS analysis of K562 cells in presence or absence of Tubastatin (TUB), Novobiocin (NOV) or Sotrastaurin (SOT) during MK differentiation (PMA) using FITC-CD41as MK differentiation marker. The graph below shows the % of differentiated cells. (B) FACS analysis of THP1 and CD34+ cells in presence or absence of Tubastatin (TUB), Novobiocin (NOV) or Sotrastaurin (SOT) during macrophage differentiation (PMA/SC-Mac) using FITC-CD68 as differentiation marker. The graph below shows the % of differentiated cells. (C) Confocal microscopy of THP1 cells undergoing macrophage differentiation (PMA) in presence or absence of Tubastatin (TUB), Novobiocin (NOV) or Sotrastaurin (SOT) showing MLLT3 (Red) distribution. Blue fluorescence (DAPI) indicates nucleus.

### MLLT3 clearance is triggered during terminal macrophage maturation, but not during intermediate monocyte differentiation

The study results from MK and macrophage differentiation prompted us to further investigate the regulation of MLLT3 during monocyte development, which serves as an intermediate differentiation stage for macrophages. To systematically evaluate this, we utilized two distinct temporal model systems. First, we employed the promyelocytic leukemia cell line HL60, which differentiates into monocytes when induced with PMA for 2 days, and further matures into macrophages when treated with PMA for 5 days (Figure 1A). Second, we utilized primary CD34+ cells cultured in monocyte-supplemented media for 14 days to induce monocyte differentiation; these same cells were subsequently cultured in macrophage-supplemented media for an additional 14 days (28 days total) to drive terminal macrophage differentiation (Figure 1A).

Immunoblot and subcellular fractionation analyses of these temporal models revealed a striking lineage-stage divergence. During the intermediate monocyte differentiation stage (2-day HL60 and 14-day CD34+ cells), total and nuclear MLLT3 levels remained entirely unchanged. Accordingly, treatment with the inhibitors TUB, NOV, or SOT had no effect on MLLT3 levels (Figure 4A, 4B & 4C). FACS and immunofluorescence analyses further confirmed that these inhibitors did not impact monocyte differentiation or alter the subcellular distribution of MLLT3 at this stage (Figure 4D & 4E).

**Figure 4.**
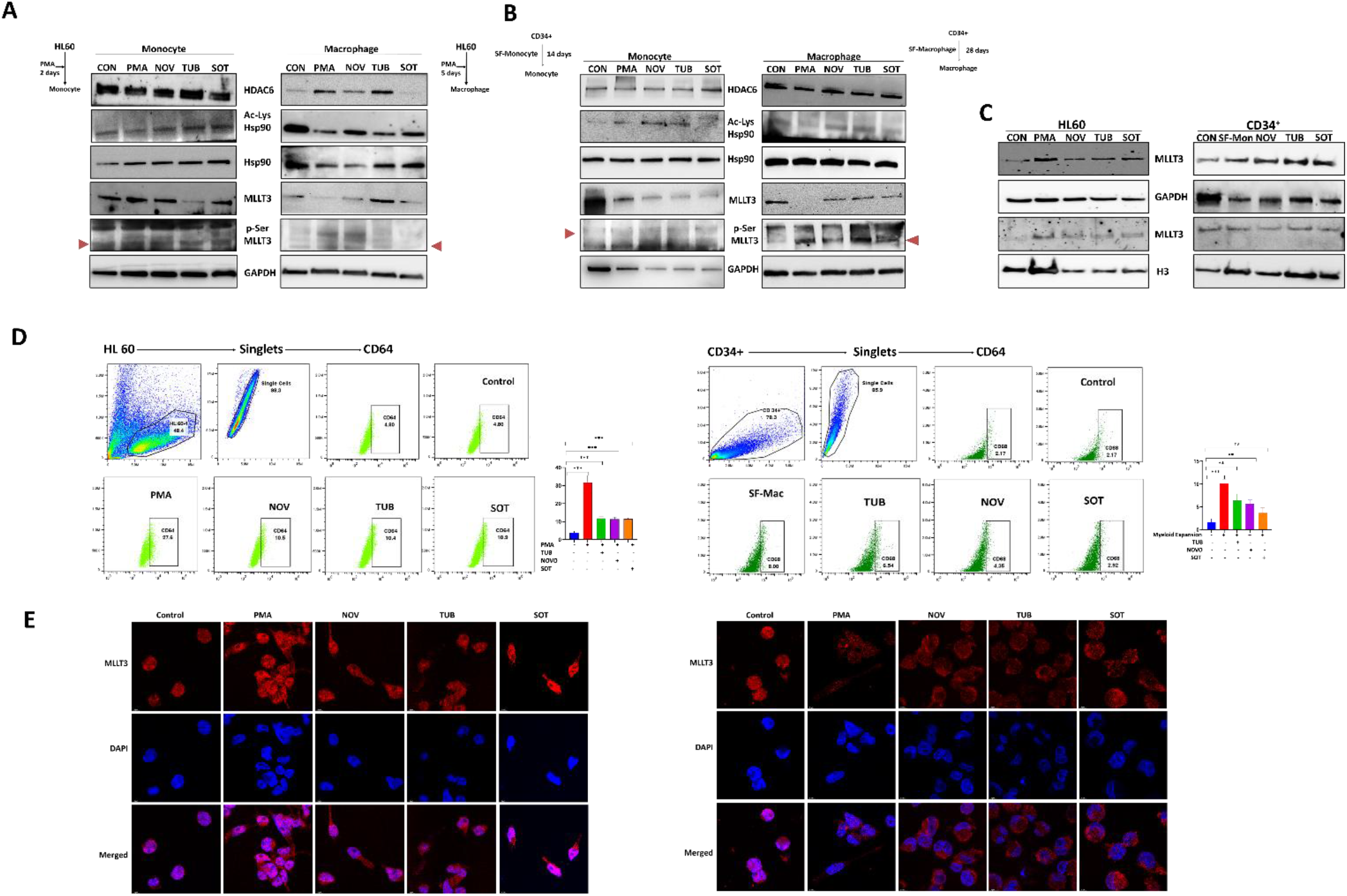
MLLT3 is not regulated by HDAC6-Hsp90 and PKC during intermediate differentiation into monocytes. (A) Immunoblot analysis of HDAC6, HSp90, Ac-Hsp90, MLLT3 and p-Ser-MLLT3 protein levels in presence or absence of Tubastatin (TUB), Novobiocin (NOV) and Sotrastaurin (SOT) during monocyte differentiation (PMA 2 days) and macrophage differentiation (PMA 5 days) of HL60 cells. (B) Immunoblot analysis of HDAC6, HSp90, Ac-Hsp90, MLLT3 and p-Ser-MLLT3 protein levels in presence or absence of Tubastatin (TUB), Novobiocin (NOV) and Sotrastaurin (SOT) during monocyte differentiation (SF-Mon) and macrophage differentiation (SF-Mac) of CD34+ cells. (C) FACS analysis of HL60 and CD34+ cells in presence or absence of Tubastatin (TUB), Novobiocin (NOV) or Sotrastaurin (SOT) during monocyte differentiation (PMA/SF-Mon) using FITC-CD64 as differentiation marker. The graph below shows the % of differentiated cells. D. Confocal microscopy of HL60 cells undergoing monocyte (PMA 2 days) and macrophage (PMA 5 days) differentiation in presence or absence of Tubastatin (TUB), Novobiocin (NOV) or Sotrastaurin (SOT) showing MLLT3 (Red) distribution. Blue fluorescence (DAPI) indicates nucleus.

**Figure 5.**
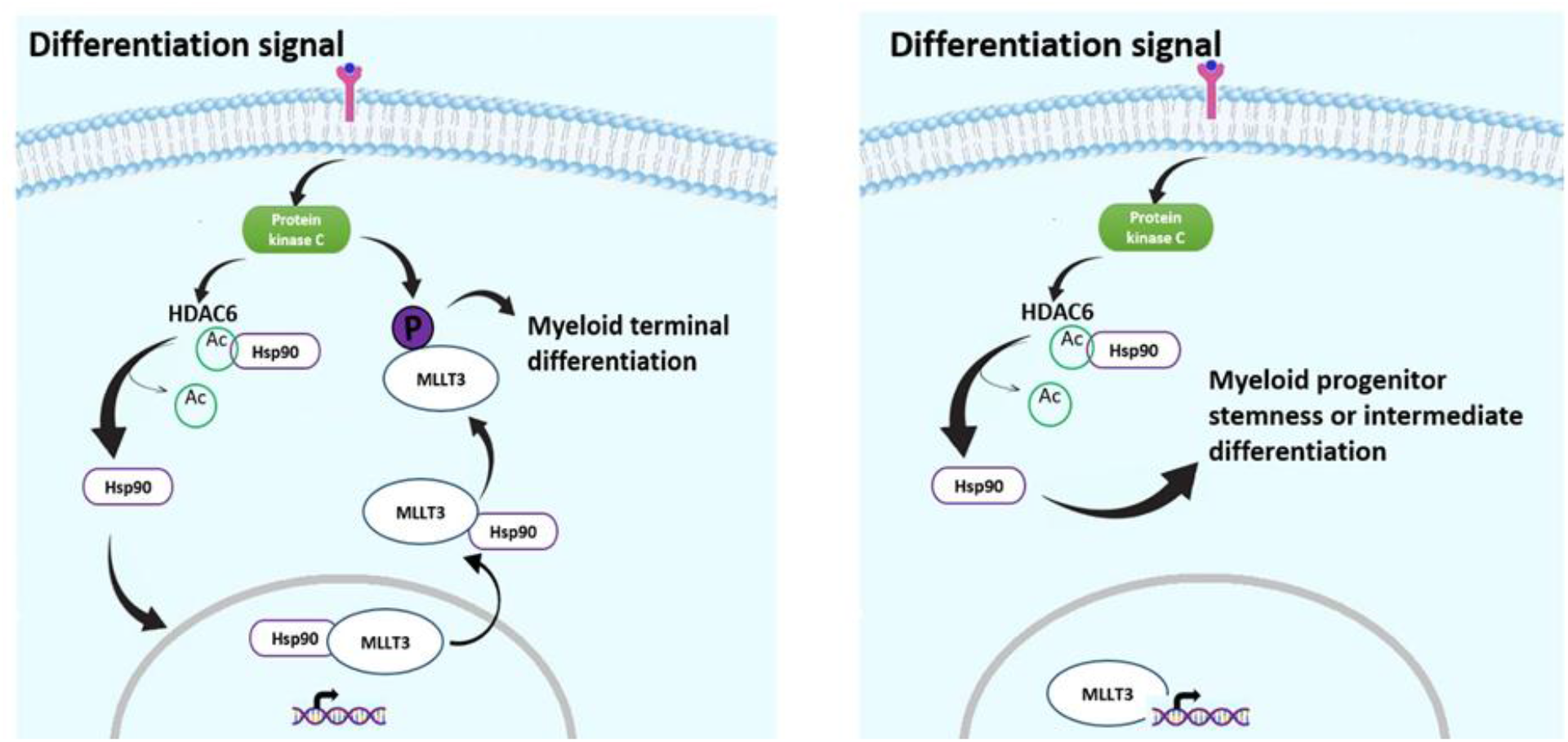
Regulation of Hsp90 (*via* deacetylation) and MLLT3 (*via* cytoplasmic localization and phosphorylation). During terminal differentiation of myeloid progenitor cells into megakaryocyte and macrophage, HSp90 undergoes deacetylation by HDAC6 and translocates MLLT3 from nucleus to cytoplasm where MLLT3 (probably) gets serine phosphorylated by PKC. But these epiproteomic changes of Hsp90 or MLLT3 does not occur during intermediate differentiation into monocytes.

In sharp contrast, progression to the terminal macrophage stage (5-day HL60 and 28-day CD34+ cells) led to a distinct decrease in MLLT3 levels. Consistent with our MK data, this decrease was successfully rescued by treatment with TUB, NOV, or SOT (Figure 4A, 4B & 4C). Furthermore, FACS analysis indicated that treatment with these inhibitors effectively blocked terminal macrophage differentiation. Immunofluorescence imaging visually confirmed the regulatory role of PKC, HDAC6, and Hsp90 on MLLT3 during this final maturation phase (Figure 4D & 4E). Together, these data demonstrate that the PKC-HDAC6-Hsp90-mediated clearance of MLLT3 is selectively activated during terminal macrophage differentiation, but remains inactive during the intermediate monocyte stage.

## Discussion

Lineage commitment and differentiation of stem cells and progenitor cells are tightly regulated by epigenetic, epitranscriptomic, and epiproteomic processes. Several proteins play crucial role in self-renewal and maintenance of stemness. MLLT3, a component of the SEC, has been identified as a key factor involved in maintaining the stemness of HSCs^3^. Despite it being associated with MLL/KMT2a in leukemogenesis, its role in the SEC remains indispensable. MLLT3 recognizes and binds to crotonylated lysines on histone tails, facilitating the release of RNA polymerase II transient pausing.^7^ Loss of MLLT3 in MEP cells leads to MK differentiation but is necessary for erythropoiesis. Nonetheless, the underlying mechanism of MLLT3 loss during MK differentiation remains unknown. Hsp90 which is a chaperone protein, has been involved in AML progression by assisting in fusion proteins refolding^19^. Hsp90 inhibitors have been explored as potential drug candidates for AML treatment^20^. Under physiological conditions, HDAC6 regulates Hsp90 chaperone activity through deacetylation^21^. In this study, we aimed to explore the loss of MLLT3 during myeloid differentiation and understand the indirect regulation of MLLT3 by HDAC6 through Hsp90 deacetylation.

Our findings, consistent with the findings of the study by Kovacs et al.^15^ demonstrated reduced nuclear levels of MLLT3 during MK differentiation. Inhibition of Hsp90 by NOV further supported this observation. Moreover, during MK differentiation, treatment with TUB showed a decrease in MLLT3 levels compared to the MLLT3 levels during treatment with NOV. Thus, indicating HDAC6’s regulatory influence over Hsp90. We then investigated the cause of MLLT3 downregulation and found evidence for serine phosphorylation-mediated degradation, as phosphorylation of serine, threonine, or tyrosine residues can lead to protein ubiquitination and degradation^22^. The data from bioinformatics tools and previous studies suggested PKC as a potential kinase responsible for MLLT3 serine phosphorylation^23^. Consistent with this, our experiments using PKC inhibitor SOT, showed decreased phosphoserine MLLT3 levels and increased levels with PMA or TPO. Thus, indicating PKC-mediated MLLT3 phosphorylation. Furthermore, reduced mRNA expression of GATA1 during MK differentiation provided evidence linking PKC-mediated MLLT3 phosphorylation to decreased nuclear abundance and subsequent MK differentiation^12^.

Our study extended the investigation to macrophage differentiation and demonstrated similar results, supporting the negative regulatory role of HDAC6 and Hsp90 over MLLT3 during MK and macrophage differentiation. However, during monocyte differentiation, MLLT3 levels remained unaffected, indicating a distinct regulatory mechanism. These findings align with earlier studies showing that MLLT3 levels did not impact granulocyte/macrophage (GM) lineage decisions in early CD34+ cell compartments, but sustained MLLT3 levels inhibited GM differentiation^12^.

In conclusion, our results support the role of HDAC6 and Hsp90 in negatively regulating MLLT3 only during terminal differentiation into MKs or macrophages, but not during intermediate monocyte differentiation. We propose that PKC-mediated serine phosphorylation of MLLT3 during terminal differentiation might be responsible for its lower nuclear abundance. Our study contributes to the growing knowledge of the intricate regulatory network governing myeloid lineage differentiation, offering potential insights for therapeutic strategies targeting hematological disorders and tissue regeneration.

Further investigations are required to address several questions: (i) The exact cellular location of MLLT3 phosphorylation-whether in the cytoplasm or nucleus, (ii) The fate of phosphorylated MLLT3-ubiquitination or cytoplasmic retention, (iii) Whether MLLT3 plays a role in cell fate decisions of the lymphoid lineage, and (iv) The involvement of HDAC6-Hsp90 in regulating MLLT3 during HSC differentiation. Addressing these challenges will provide a comprehensive understanding of the regulation of MLLT3 and its implications in HSC lineage commitment.

## Data availability

No datasets were generated or analyzed during the current study.

## References

1. Biswas, A. & Hutchins, R. Embryonic stem cells. Stem Cells Dev 16, 213–22 (2007).

2. Eaves, C. J. Hematopoietic stem cells: concepts, definitions, and the new reality. Blood 125, 2605–13 (2015).

3. Calvanese, V. et al. MLLT3 governs human haematopoietic stem-cell self-renewal and engraftment. Nature 576, 281–286 (2019).

4. Lin, C. et al. AFF4, a component of the ELL/P-TEFb elongation complex and a shared subunit of MLL chimeras, can link transcription elongation to leukemia. Mol Cell 37, 429–37 (2010).

5. He, N. et al. HIV-1 Tat and host AFF4 recruit two transcription elongation factors into a bifunctional complex for coordinated activation of HIV-1 transcription. Mol Cell 38, 428–38 (2010).

6. Li, Y. et al. AF9 YEATS domain links histone acetylation to DOT1L-mediated H3K79 methylation. Cell 159, 558–71 (2014).

7. Li, Y. et al. Molecular Coupling of Histone Crotonylation and Active Transcription by AF9 YEATS Domain. Mol Cell 62, 181–193 (2016).

8. Zhang, Q. et al. Structural Insights into Histone Crotonyl-Lysine Recognition by the AF9 YEATS Domain. Structure 24, 1606–12 (2016).

9. Schoch, C. et al. AML with 11q23/MLL abnormalities as defined by the WHO classification: incidence, partner chromosomes, FAB subtype, age distribution, and prognostic impact in an unselected series of 1897 cytogenetically analyzed AML cases. Blood 102, 2395–402 (2003).

10. Krivtsov, A. V et al. Transformation from committed progenitor to leukaemia stem cell initiated by MLL-AF9. Nature 442, 818–22 (2006).

11. Pina, C., May, G., Soneji, S., Hong, D. & Enver, T. MLLT3 regulates early human erythroid and megakaryocytic cell fate. Cell Stem Cell 2, 264–73 (2008).

12. Jackson, S. E. Hsp90: structure and function. Top Curr Chem 328, 155–240 (2013).

13. Truman, A. W. et al. Expressed in the yeast Saccharomyces cerevisiae, human ERK5 is a client of the Hsp90 chaperone that complements loss of the Slt2p (Mpk1p) cell integrity stress-activated protein kinase. Eukaryot Cell 5, 1914–24 (2006).

14. Lin, J. J. & Hemenway, C. S. Hsp90 directly modulates the spatial distribution of AF9/MLLT3 and affects target gene expression. J Biol Chem 285, 11966–73 (2010).

15. Kovacs, J. J. et al. HDAC6 regulates Hsp90 acetylation and chaperone-dependent activation of glucocorticoid receptor. Mol Cell 18, 601–7 (2005).

16. Messaoudi, K. et al. Critical role of the HDAC6-cortactin axis in human megakaryocyte maturation leading to a proplatelet-formation defect. Nat Commun 8, 1786 (2017).

17. Doddi, S. K., Kummari, G. M.V. J. & Kalle, A. M. Protein kinase A mediates novel serine-584 phosphorylation of HDAC4. Biochemistry and Cell Biology 97, 526–535 (2019).

18. Vanaja, G. R., Ramulu, H. G. & Kalle, A. M. Overexpressed HDAC8 in cervical cancer cells shows functional redundancy of tubulin deacetylation with HDAC6. Cell Communication and Signaling 16, 20 (2018).

19. Flandrin, P. et al. Significance of heat-shock protein (HSP) 90 expression in acute myeloid leukemia cells. Cell Stress Chaperones 13, 357–64 (2008).

20. Lazenby, M., Hills, R., Burnett, A. K. & Zabkiewicz, J. The HSP90 inhibitor ganetespib: A potential effective agent for Acute Myeloid Leukemia in combination with cytarabine. Leuk Res 39, 617–24 (2015).

21. Du, Y. et al. HDAC6-mediated Hsp90 deacetylation reduces aggregation and toxicity of the protein alpha-synuclein by regulating chaperone-mediated autophagy. Neurochem Int 149, 105141 (2021).

22. Filipčík, P., Curry, J. R. & Mace, P. D. When Worlds Collide-Mechanisms at the Interface between Phosphorylation and Ubiquitination. J Mol Biol 429, 1097–1113 (2017).

23. Zhang, W. et al. Aldosterone-induced Sgk1 relieves Dot1a-Af9-mediated transcriptional repression of epithelial Na+ channel alpha. J Clin Invest 117, 773–83 (2007).

